# NSUN2 introduces 5-methylcytosines in mammalian mitochondrial tRNAs

**DOI:** 10.1101/626960

**Authors:** Lindsey Van Haute, Song-Yi Lee, Beverly J. McCann, Christopher A. Powell, Dhiru Bansal, Caterina Garone, Sanghee Shin, Jong-Seo Kim, Michaela Frye, Joseph G. Gleeson, Eric Miska, Hyun-Woo Rhee, Michal Minczuk

**Affiliations:** Medical Research Council Mitochondrial Biology Unit, University of Cambridge, Cambridge, CB2 0XY, UK; Department of Chemistry, Seoul National University, Gwanak-ro 1, Seoul 08826, South Korea; Wellcome Trust/Cancer Research UK Gurdon Institute, University of Cambridge, Tennis Court Road, Cambridge CB2 1QN, UK; Department of Genetics, University of Cambridge, Downing Street, Cambridge CB2 3EH, UK; Center for RNA Research, Institute of Basic Science, Seoul 08826, Korea; School of Biological Sciences, Seoul National University, Seoul 08826, Korea; German Cancer Center (DKFZ), Im Neuenheimer Feld 280, 69120 Heidelberg, Germany; Rady Children’s Institute for Genomic Medicine, Rady Children’s Hospital, San Diego, CA, 92123, USA; Wellcome Sanger Institute, Wellcome Trust Genome Campus, Cambridge CB10 1SA, UK

## Abstract

Maintenance and expression of mitochondrial DNA is indispensable for proper function of the oxidative phosphorylation machinery. Post-transcriptional modification of mitochondrial RNA has emerged as one of the key regulatory steps of human mitochondrial gene expression. Mammalian NOP2/Sun RNA Methyltransferase Family Member 2 (NSUN2) has been characterised as an RNA methyltransferase that introduces 5-methylcytosine (m^5^C) in nuclear-encoded tRNAs, mRNAs, microRNA and noncoding RNAs. In these roles, NSUN2 has been associated with cell proliferation and differentiation. Pathogenic variants in NSUN2 have been linked with neurodevelopmental disorders. Here we employ spatially restricted proximity labelling and immunodetection to demonstrate that NSUN2 is imported into the matrix of mammalian mitochondria. Using three genetic models for NSUN2 inactivation – knockout mice, patient-derived fibroblasts and CRISPR/Cas9 knockout in human cells – we show that NSUN2 in necessary for the generation of m^5^C at positions 48, 49 and 50 of several mammalian mitochondrial tRNAs. Finally, we show that inactivation of NSUN2 does not have a profound effect on mitochondrial tRNA stability and oxidative phosphorylation in differentiated cells. We discuss the importance of the newly discovered function of NSUN2 in the context of human disease.

## Introduction

Mammalian mitochondria contain small semi-independent genomes referred to as mtDNA, which encode the key subunits of the oxidative phosphorylation (OXPHOS) system. The concerted action of both the nuclear and mitochondrial genomes is required for OXPHOS and the synthesis of adenosine triphosphate (ATP). MtDNA codes for 13 proteins, 2 mitochondrial (mt-) rRNA and 22 mt-tRNA, required for intramitochondrial protein synthesis. These combine with ∼90 nuclear-encoded proteins to form five OXPHOS enzyme complexes. An additional 200-300 mammalian genes are involved in mtDNA maintenance, in various steps of mitochondrial protein synthesis, and assembly of the respiratory chain complexes.

Universally, RNA molecules undergo multiple post-transcriptional modifications, which are crucial for their biogenesis and/or function. The mitochondrially-encoded tRNAs also undergo extensive post-transcriptional maturation including chemical nucleotide modifications, CCA addition at the 3’ end and deadenylation (1,2). Recent data suggest that over 7% of all mt-tRNA residues in mammals undergo post-transcriptional modification, with over 30 different modified mt-tRNA positions so far described (3). The nucleotide modifications of mt-tRNA play a role in decoding, tRNA structural stability, and aminoacylation (4).

Cytosine-5 methylation is a common epigenetic modification found in DNA where it plays important regulatory roles in transcription (5). It is also the most abundant post-transcriptional RNA modification (6) and several enzymes responsible for these RNA modifications have been identified. With the exception of DNA methyltransferase 2 (Dnmt2), which catalyses m^5^C formation of position C38 of cytoplasmic (cyto-) tRNA^Asp^, tRNA^Val^ and tRNA^Gly^ (7,8), all known RNA methyltransferases are members of the highly conserved NOL1/NOP2/Sun (NSUN) domain containing family. NSUN1 and NSUN5 are involved in m^5^C modification of rRNAs for cytoplasmic ribosomes (9,10), while cytoplasmic tRNA^Cys^ and tRNA^Thr^ (position 72) are the substrates of NSUN6 (11). Two NSUN family members are imported and function in mitochondria. NSUN4 methylates mitochondrial 12S mt-rRNA and plays a role in mitochondrial ribosome biogenesis (12). Furthermore, our group and others have demonstrated that NSUN3 is responsible for methylation of position C34 of mt-tRNA^Met^ (13-15), which is further oxidised to 5-formylcytosine by AlkBH1 (15,16). NSUN2 is a well-characterized RNA methyltransferase, responsible for cytosine-5 methylation (m^5^C) of tRNA^Leu(CAA)^ at position C34 and C40 of tRNA^Gly^, at C48 and 49 of tRNA^Asp^, tRNA^Val^ and tRNA^Gly^ and at C50 of tRNA^Gly^ (17,18). In addition to cytoplasmic tRNAs this protein also methylates a variety of cytoplasmic mRNAs (19) and vault non-coding RNAs that can function as microRNAs (20).

Recent data indicate that defects in the nucleotide modification of mt-tRNA can lead to human disorders of mitochondrial respiration (1,21-23). The m^5^C modification in the variable-loop (V-loop) region has been detected in bovine and human mt-RNAs (4). However, the biogenesis of this mitochondrial m^5^C has not been investigated. Here we demonstrate that NSUN2, which previously has only been associated with the modification of extramitochondrial RNA, is partially localised in the mitochondrial matrix and is responsible for methylation of tRNA positions 48, 49 and 50 in the V-loop region of six human and eight mouse mt-tRNA. Although ablation of NSUN2 results in undetectable levels of mt-tRNA m^5^C, the lack of this modification does not appear to have any substantial effect on mammalian mitochondrial function. Taken together, this work demonstrates that, in addition to its role in the modification of nuclear-encoded RNA, NSUN2 is also responsible for the majority of m^5^C modifications of mt-tRNAs in mouse and human.

## Materials and Methods

### Animals and housing

All animal experiments were carried out in accordance with the UK Animals (Scientific Procedures) Act 1986 (UK Home Office license: PPL70/7538, PPL80/2231 and PPL80/2619) and EU Directive 2010/63/EU. The NSUN2 knockout mice generated as described previously (24) were housed in the Wellcome Trust - Medical Research Council Cambridge Stem Cell Institute Animal Unit.

### Maintenance and transfection of mammalian cell lines

All cells were grown at 37°C in a humidified atmosphere with 5% CO2 in high-glucose DMEM supplemented with 10% foetal bovine serum. The Flp-In T-Rex HEK293T cell line was used to generate a doxycycline-inducible expression of NSUN2.FLAG.STREP2. The full-length NSUN2 cDNA construct was cloned into a pcDNA5/FRT/TO plasmid encoding FLAG.STREP2-tag as previously described (25). The oligonucleotides used for cloning can be found in Supplementary Table S1. Cells were grown in the same medium described above, supplemented with selective antibiotics hygromycin (100 µg/mL) and blasticidin (15 µg/mL). For transient complementation of U2OS NSUN2 KO cells were transfected with 0.8 µg of NSUN2.FLAG.STREP2 construct with Lipofectamin 2000. Medium was replaced after 24h and cells were analysed after 48h.

### Genome editing by CRISPR/Cas9

To generate NSUN2 knockout in human cells, a pair of gRNAs against NSUN2 (**Supplementary Table S1**) were cloned in All-in-One-GFP plasmid as described in Chiang *et al*. (26). The plasmid was transfected into human osteosarcoma U-2 (U2OS) cells by electroporation using the Neon Transfection System (Life Technologies) according to the manufacturer’s instructions. 24 hours after transfection, cells were FACS-sorted based on EGFP expression and plated to obtain individual clones. NSUN2 KO clones were analysed by PCR and Sanger sequencing to detect NHEJ-related nuclear genome edits, RT-qPCR (using SYBR Green Master Mix with 100 ng cDNA input) to analyse steady-state levels of NSUN2 mRNA (**Supplementary Table S2**) and Western blotting to examine NSUN2 protein levels (**Supplementary Figure S1**).

### Spatially restricted proximity labelling and MS analysis

For subcellular proteome mapping, we prepared several HEK293 cell lines stably expressing various sub-mitochondrially localised constructs of APEX2. The specific sub-mitochondrial localisation was achieved by fusing APEX2 to a well-characterised targeting signal directing it to the mitochondrial matrix (MLATRVFSLVGKRAISTSVCVRAH from cytochrome c oxidase subunit 4 isoform 1 (27), Mito-APEX2), a protein localised in intermembrane space - SCO1 (IMM-APEX2), or a protein localised in outer mitochondrial membrane - TOM20 (OMM-APEX2). The cytosol-localised APEX2 (Cyto-APEX2) was included as additional control. These constructs have been described previously (28). We have cultured HEK293 cells in T75 flask and expression of genetically targeted sub-mitochondrial APEX2 (matrix, IMS-, OMM-APEX2) and APEX2-NES was induced by doxycycline (100 ng/ml) overnight. Next, we incubated the cells with desthiobiotin-phenol (DBP, 250 µM) for 30 min at 37 °C, followed by 1 min incubation with 1 mM H_2_O_2_ to generate the DBP radical *in situ* (29). After labelling, the cells were lysed with 2% SDS in 1x TBS (25 mM Tris, 0.15 M NaCl, pH 7.2) lysis buffer and DBP-labelled peptides were enriched by streptavidin and analyzed by LC-MS/MS.

Analytical capillary columns (100 cm × 75 µm i.d.) and trap columns (2 cm × 150 µm i.d) were packed in-house with 3 µm Jupiter C18 particles (Phenomenex, Torrance, CA). The long analytical column was placed in a column heater (Analytical Sales and Services, Pompton Plains, NJ) regulated to a temperature of 45 °C. Dionex Ultimate 3000 RSLC nano-system (Thermo Scientific, Sunnyvale, CA) was operated at a flow rate of 350 nL/min over 2 hours with linear gradient ranging from 95 % solvent A (H_2_O with 0.1% formic acid) to 40% of solvent B (acetonitrile with 0.1% formic acid). The enriched samples were analysed on an Orbitrap Fusion Lumos mass spectrometer (Thermo Scientific) equipped with an in-house customized nanoelectrospray ion source. Precursor ions were acquired (m/z 300 – 1500 at 120 K resolving power and the isolation of precursor for MS/MS analysis was performed with a 1.4 Th. Higher-energy collisional dissociation (HCD) with 30% collision energy was used for sequencing with a target value of 1e5 ions determined by automatic gain control. Resolving power for acquired MS2 spectra was set to 30 k at m/z 200 with 150 ms maximum injection time.

All MS/MS data were searched by MaxQuant (version 1.5.3.30) with Andromeda search engine at 10 ppm precursor ion mass tolerance against the SwissProt Homo sapiens proteome database (20199 entries, UniProt (http://www.uniprot.org/)). The following search parameters were applied: trypic digestion, fixed carbaminomethylation on cysteine, dynamic oxidation of methionine, DBP (331.190) labels of tyrosine residue. The False discovery rate (FDR) was set at <1% for peptide spectrum match including unlabelled peptides. FDR less than 1% was obtained for unique labelled peptide and unique labelled protein level as well.

### Cell fractionation

HEK293T cells were cultured in DMEM supplemented with 10% FBS. Detached cells were washed twice with PBS and resuspended in homogenisation buffer containing mannitol (0.6 M), Tris (10 mM), EDTA (0.1 mM) (pH 7.4) and protease inhibitors (Roche). Cells were homogenized with a glass Dounce homogenizer. Next, differential centrifugation was used to obtain the different cell fractions (1000 x g for 10 min to remove cell debris and nuclei, next for 10 min at 10000 x g to pellet the mitochondria). The pellet containing the mitochondrial fractions was resuspended in 50 mM TrisHCl, pH 7.4 with 150 mM NaCl and divided in four fractions of which three were used for treatment with Proteinase K. Proteinase K treatment was performed under different stringencies: 0,2 µg / 5 mg for 10 min, 2 µg / 5 mg for 20 min and 20 µg per 5 mg mitochondrial proteins for 30 min.

### Immunodetection of proteins

The intracellular localization of NSUN2 by immunofluorescence in fixed human osteosarcoma 143B cells was performed as described previously (30). The following antibodies were used: rabbit anti-TOM20 (Santa Cruz Biotechnology, sc-1145, 1:200), Rat anti-HA (Roche, 118667431001, 1:200), Alexa Fluor 594 anti-rabbit (1:1000, ab150088, Abcam), Alexa Fluor 488 anti rat (Abcam, ab150157, 1:1000), mouse anti-FLAG (Sigma, F1084, 1:500). Immunofluoresence images were captured using a Zeiss LSM880 confocal microscope and processed using ImageJ.

For western blot analysis, 10-30 µg of extracted proteins were loaded on SDS-PAGE 4-12% bis-tris gels (Life Technologies) and transferred onto a membrane using iBlot 2 Dry Blotting System (Thermo Fisher Scientific). The following antibodies were used: rabbit anti-NSUN2 (Proteintech, 20854-1-AP, 1:1000), mouse anti-Tubulin (Sigma Aldrich, T9026, 1:5000), mouse anti-TOM22 (Abcam, ab10436, 1:2000), mouse anti-GAPDH (Abcam, ab9484, 1:5000) mouse anti-FLAG (Sigma, F3165, 1:1000), mouse anti-mtSSB1 (a kind gift of Prof. D. Kang, 1:4000), total OXPHOS Mouse WB antibody cocktail (Abcam, ab110412, 1:2000) and mouse anti-HSC70 (Santa Cruz Biotechnology sc7298, 1:10000).

### Mitochondrial DNA copy number analysis

Total cellular DNA was isolated from mouse liver and heart tissue. Tissue was cut into little pieces in medium (pH 7.4) containing 0.32M sucrose, 10mM Tris-HCl pH 7.4 and 1 mM EDTA and homogenized. DNA was extracted using a DNeasy Blood and Tissue kit (Qiagen) according to the manufacturer’s recommendations. Copy numbers of mtDNA were measured by quantitative PCR according to the manufacturer’s recommendations and using PowerUp SYBR Green Mister Mix (Applied Biosystems). Primer and probe sequences can be found in **Supplementary Table S2**.

### RNA analysis by northern blot

RNA was extracted with TRIzol reagent (Thermo Fisher Scientific). Total RNA was resolved on 1% agarose gels containing 0.7?M formaldehyde in 1 × MOPS buffer (mt-mRNAs) or on 10% UREA–PAGE (mt-tRNAs), transferred to a nylon membrane, ultraviolet-crosslinked to the membrane and hybridized with radioactively labelled T7-transcribed radioactive RNA-probes.

### RNA bisulfite sequencing

RNA was extracted with TRIzol reagent (Thermo Fisher Scientific) and treated with TURBO DNase (Ambion) according to manufacturer’s instructions. Bisulfite conversion of 2 µg DNase treated RNA was performed using the Methylamp One-Step DNA modification kit (Epigentek). The reaction mixture was incubated for three cycles of 90°C for 5 min and 60°C for 1h. Following desalting with Micro Bio-spin 6 chromatography columns (Bio-Rad) twice, RNA was desulfonated by adding an equal volume of 1M Tris (pH 9.0) and incubated for 1h at 37°C. For whole genome sequencing of mouse RNA a library was generated with Truseq Small RNA Library Preparation kit (Illumina) according to the manufacturer’s protocol. Alternatively, reverse transcription was performed with specific primers (Superscript III, Life technologies). After the first stage PCR with overhang primers (**Supplementary Table S2**), excess primers were removed with Ampure XP beads. An 8 cycles second round PCR was performed with indexed primers (Nextera XT), followed by another clean-up with Ampure XP beads. Quality and concentration were assessed with a D1000 Screentape for TapeStation (Agilent Genomics). Libraries were subjected to high-throughput sequencing using the Illumina MiSeq platform. After quality trimming and 3’ end adaptor clipping with Trim_Galore!, reads longer than 20 nt were aligned to a computationally bisulfite-converted human reference genome (GRCh38) with Bismark (31).

## Results

### A proportion of NSUN2 localises inside human mitochondria

In order to obtain data on the subcellular localisation of endogenous NSUN2 in human cells, we conducted spatially restricted proximity labelling experiments (32). We targeted APEX2 to four different subcellular compartments of HEK293 cells: mitochondrial matrix (using the mitochondrial targeting sequence of cytochrome c oxidase fused to APEX2: Matrix-APEX2), the mitochondrial intermembrane space (using SCO1 fused to APEX2: IMS-APEX2), outer-mitochondrial membrane (using TOM20 fused to APEX2: OMM-APEX2) and the cytosol (using a nuclear export signal, NES, fused to APEX2: Cyto-APEX2) (**Figure 1A**)(28). Within these subcellular localisations, we found that endogenous NSUN2 was reproducibly labelled by matrix-APEX2, while other APEX2 constructs showed no labelling on this protein (**Figure 1B**). Direct mass spectrometric (MS) analysis detected the desthiobiotin-phenol (DBP)-labelled tyrosine site (Y646) of NSUN2 in the Matrix-APEX2 experiment (**Figure 1C**). Given that the *in situ* generated phenoxyl radical by Matrix-APEX2 cannot cross the inner-mitochondrial membrane due to its short lifetime, this latter result suggests a mitochondrial matrix localisation of endogenous NSUN2 (29,33).

**Figure 1.**
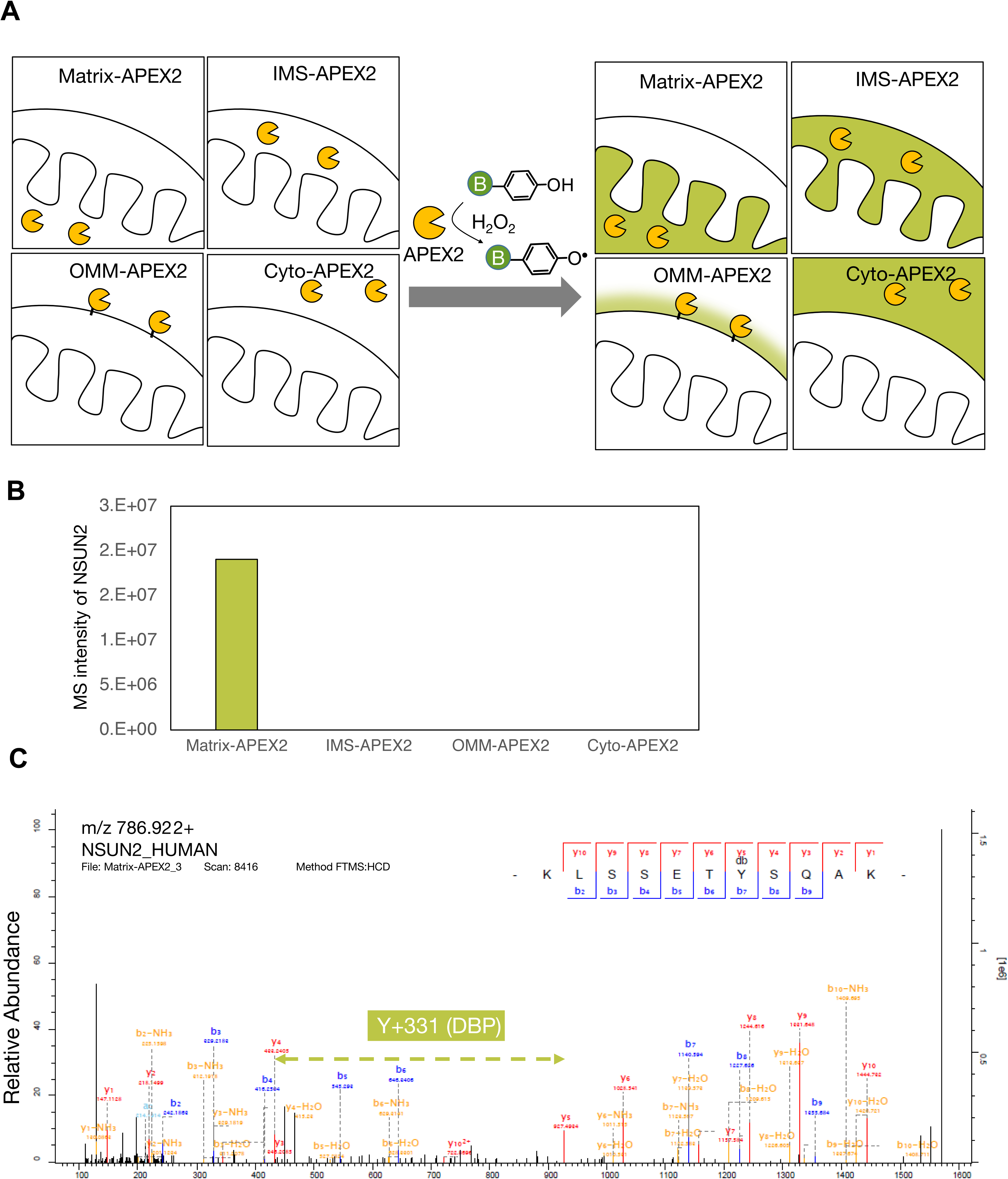
Proximity labelling reveals mitochondrial matrix pool of NSUN2. (**A**) Scheme of compartment-specific APEX2 labelling (Matrix, IMS, OMM and cytosol) (**B**) Mass spectroscopy signal intensity of APEX2-labeled NSUN2 peptides from four subcellular APEX2 constructs. (**C**) MS/MS spectra of NSUN2 DBP-peptide (KLSSET<u>Y</u>SQAK) labelled by mitochondrial matrix-targeted APEX2.

In order to provide further evidence for mitochondrial localisation of NSUN2, an HA-tagged version of the NSUN2 protein (GenBank: NM_017755.6) was transiently expressed in human osteosarcoma cells. Immunofluorescence analysis showed a variation in the cellular localisation of the NSUN2 protein. In a predominant subset of cells, NSUN2 could be detected in the nucleus or in the nucleus and cytoplasm (**Figure 2A**), consistent with previously published data (34). However, a subgroup of cells existed with predominant cytoplasmic localisation of NSUN2, in which a proportion of NSUN2 co-localised with the mitochondrial protein TOM20 (**Figure 2B**, inset). To provide additional confirmation for mitochondrial localisation of the NSUN2 protein, we performed cellular fractionation experiments of HEK293T cells inducibly expressing the NSUN2.FLAG.STREP2 construct followed by detection of the expressed protein by western blotting (**Figure 2C**). A proportion of the NSUN2.FLAG.STREP2 protein was detected in the mitochondrial fraction, along with a known mitochondrial matrix protein, mtSSB1. Both NSUN2 and mtSSB1 were resistant to proteinase K treatment of the mitochondrial fraction, consistent with NSUN2 being present within the mitochondria in human cells. Taken together, these results imply that, in addition to the cell nucleus, a proportion of NSUN2 localises inside mitochondria.

**Figure 2.**
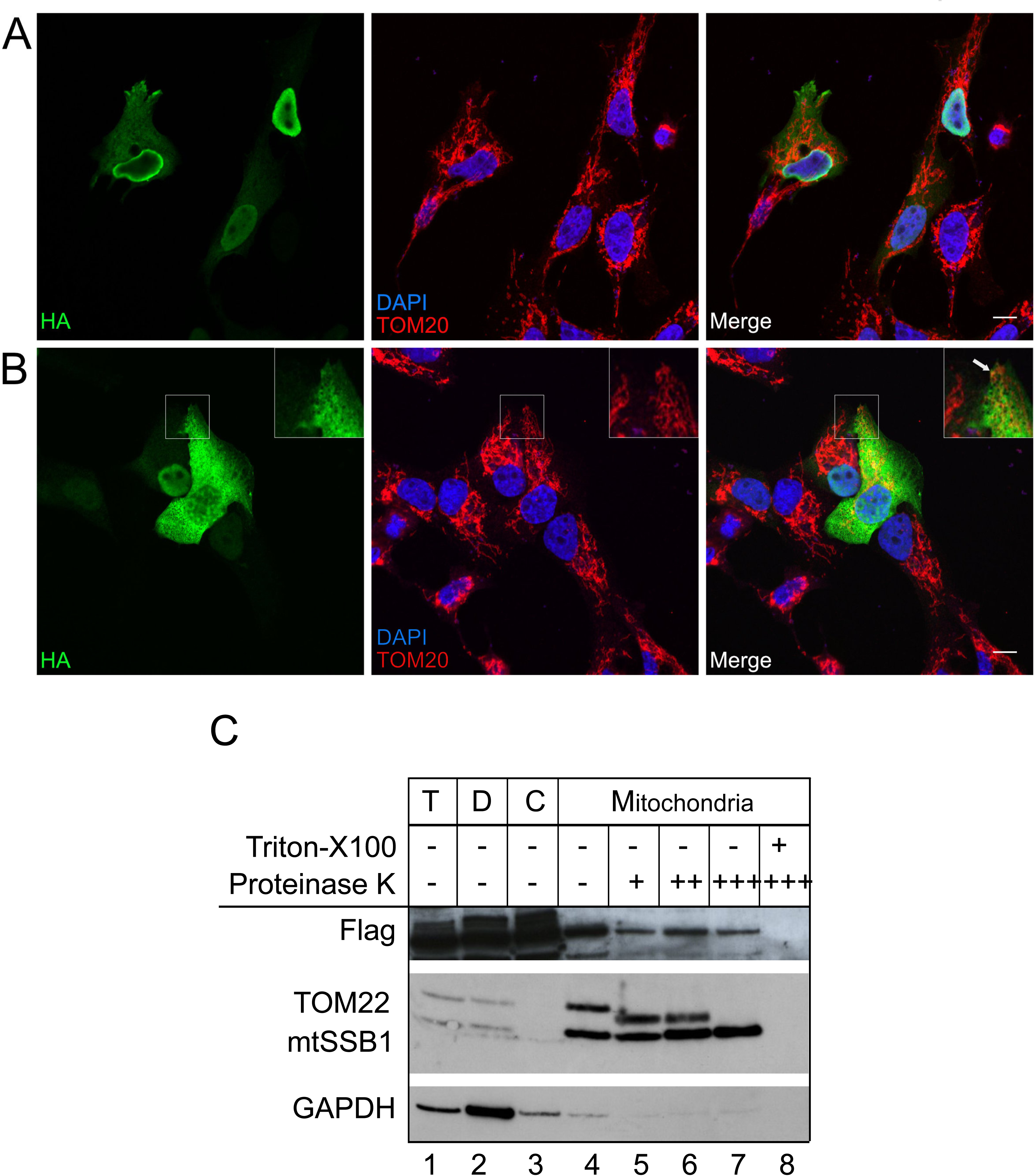
A subset of NSUN2 protein localizes in the mitochondria. (**A-B**) Immunofluorescence labelling of a HA-tagged NSUN2 construct (green) in osteosarcoma 143B cells. Cells were counterstained for the mitochondrial import receptor subunit TOM20 (red) and DAPI (blue). Scale bar: 10µM. NSUN2 localizes mainly to the nucleus (A). A fraction of the NSUN2 protein co-localizes with the mitochondria (B). (**C**) Sub-cellular localization of a FLAG-tagged NSUN2 construct analysed by western blotting with antibodies against FLAG, TOM22 (mitochondrial outer membrane), mtSSB1 (mitochondrial matrix) and GAPDH (cytosol). HEK293T cells were fractionated into debris (“D”), cytosol (“C”) and mitochondria. “T” indicates the total cell lysate.

### Mouse NSUN2 methylates mitochondrial tRNA at positions 48-50

Having shown mitochondrial localisation of the NSUN2 protein, we set out to determine whether inactivation of its expression affects m^5^C methylation of mtRNA. To this end we analysed m^5^C methylation in mouse mtRNA isolated from NSUN2-null mice (NSUN2 -/-) (24), or control animals, using high-throughput sequencing of cDNA libraries obtained after bisulfite treatment of RNA. The bisulfite RNA sequencing (BS RNA-Seq) analysis of total RNA isolated from the liver of NSUN2 -/- mice and age-matched wild type mice revealed a strong, general reduction of the m^5^C signal for mtRNA in the NSUN2 mutants (**Figure 3A**). More detailed inspection revealed that m^5^C signal reduction in the NSUN2 -/- samples corresponded to the mt-tRNA-specific reads, but not to the mt-mRNA sequences (**Figure 1B**). We next examined the total mt-tRNA pool for the tRNA positions affected by the lack of NSUN2. The methylation of the variable loop (V-loop), in particular positions 48, 49 and 50, was reduced, while the methylation within the anticodon loop, remained unaffected (**Figure 3C**). Since some mt-tRNAs were insufficiently covered in this dataset, we also used targeted resequencing of bisulfite treated RNA samples to examine whether NSUN2 is responsible for methylation of mt-tRNAs that have at a cytosine in position 48, 49 or 50. First we analysed mouse mt-tRNA^Met^ which contains two m^5^C modifications, positions C34 (wobble base, mtDNA position m.3875) and C47 (V-loop, mtDNA position m.3888) (**Figure 4A**). The BS RNA-Seq analysis revealed that the mt-tRNA^Met^ wobble base C34 is not affected by the lack of NSUN2, consistent with the previous reports on NSUN3 being responsible for modifying this base (13-15). On the contrary, methylation levels of position C47, which is almost fully methylated in wild-type mouse tissue, were reduced to background levels in the NSUN2 -/- mice samples (**Figure 4B-C**). Next, we studied m^5^C methylation in other mouse mt-tRNAs that contain cytosines in the V-loop region, namely mt-tRNA^Leu(UUR)^ (C48), mt-tRNA^His^ (C48), mt-tRNA^Glu^ (C49), mt-tRNA^Asn^ (C48), mt-tRNA^Tyr^ (C48), mt-tRNA^Leu(CUN)^ (C48 and C49) and mt-tRNA^Ser(AGY)^ (C49 and C50) (**Figure 4D-I**). The BS RNA-Seq analysis confirmed that cytosines at position 48 or 49 in mt-tRNA^Leu(UUR)^, mt-tRNA^His^, mt-tRNA^Glu^, mt-tRNA^Asn^ or mt-tRNA^Tyr^ are methylated by NSUN2 (**Figure 4D-E**). In the case of murine mt-tRNA^Leu(CUN)^, which contains two cytosines in the V-loop region (C48 and C49), position 48 and 49 are methylated in a NSUN2-dependent manner to approximately 5% and 50%, respectively, (**Figure 4F-G**). Also, in the case of murine mt-tRNA^Ser(AGY)^ which contains two cytosines in the V-loop region (C49 and C50) only position 49 is substantially methylated by NSUN2 (**Figure 4H-I**). This is somewhat similar to the m^5^C profile detected for the bovine tRNA^Ser(AGY)^ which contains three cytosines in the V-loop region (C48-50), with only C49 being m^5^C modified. Taken together these data show that ablation of mouse NSUN2 results in substantial loss of m^5^C in the V-loop region of eight mt-tRNAs, consistent with this protein being responsible for the introduction of this modification.

**Figure 3.**
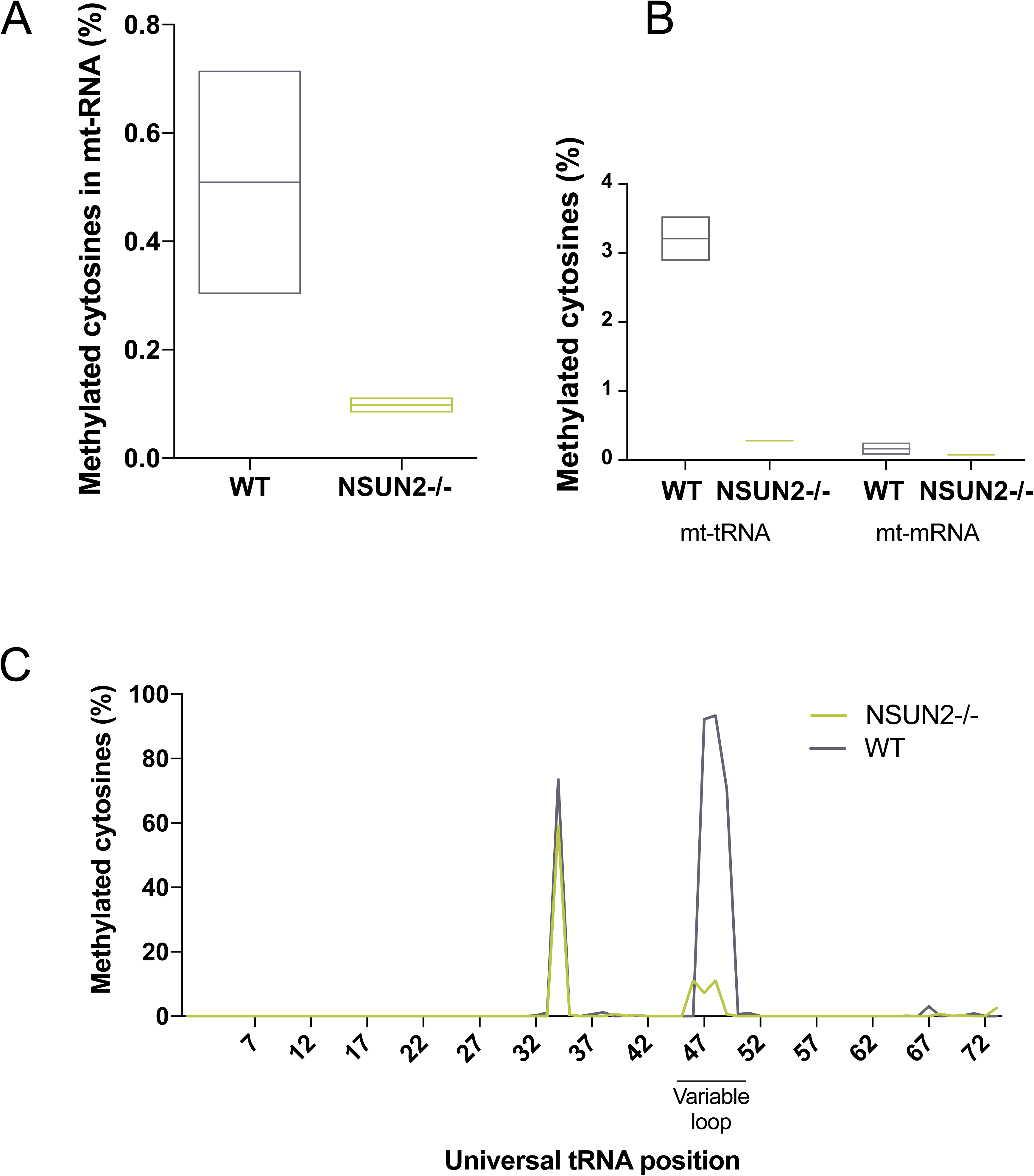
Lack of NSUN2 in mouse affects mt-RNA cytosine methylation. (**A-C**) Whole genome sequencing of bisulfite treated RNA isolated from liver tissue from NSUN2 -/- and WT mice. (**A**) Total mitochondrial RNA cytosine methylation in WT and NSUN2 -/- mouse liver tissue. (**B**) Cytosine methylation of mitochondrial tRNA and mRNA in WT and NSUN2 -/- mice. (**C**) Cytosine methylation level of the mt-tRNA pool for each position of the mt-tRNA (according to the universal tRNA position) for WT and NSUN2 -/- mice.

**Figure 4.**
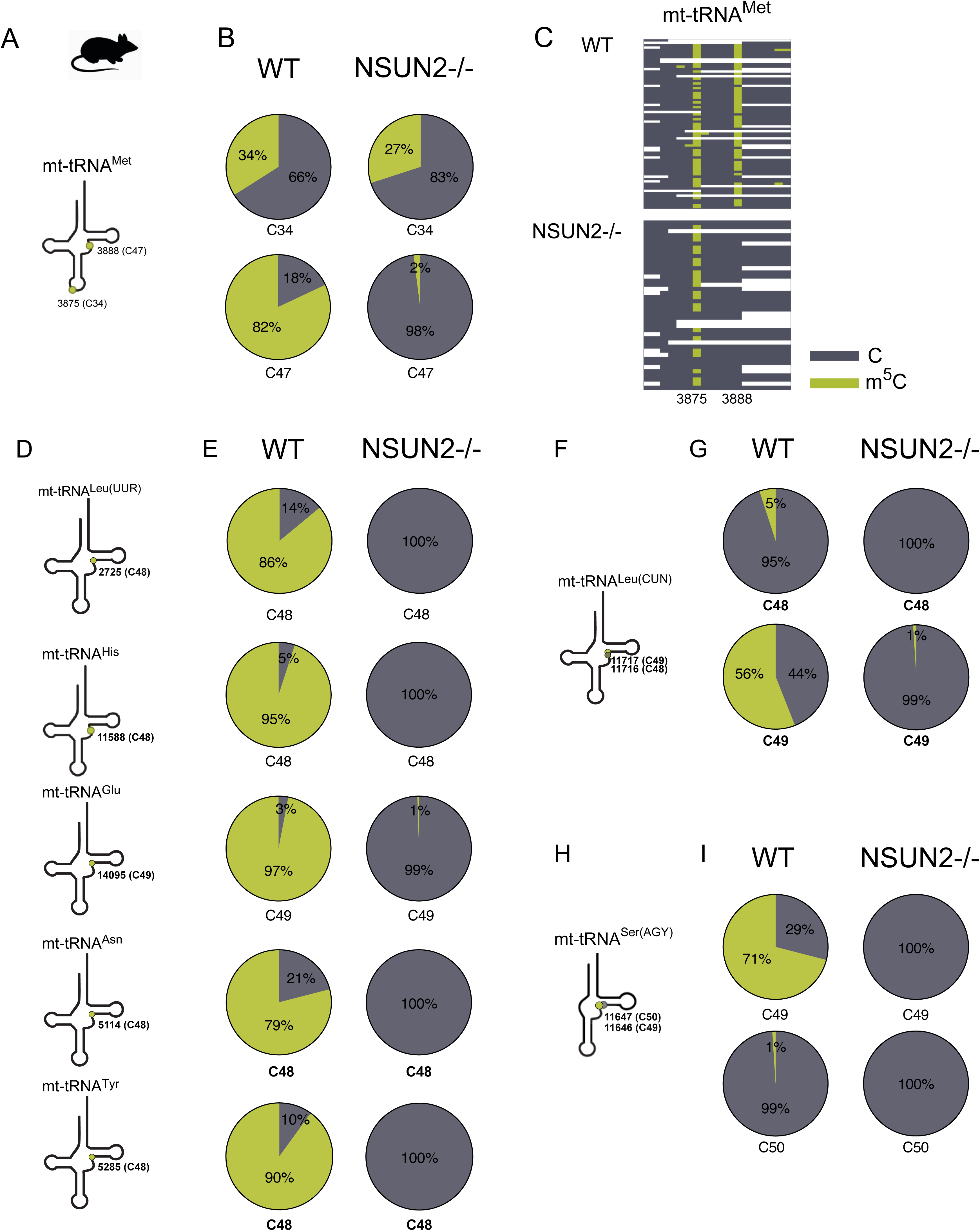
NSUN2 methylates positions C48-49 of mitochondrial tRNA in mouse. (**A-I**) Cytosine methylation of mouse liver mt-tRNAs. Throughout the figure green indicates methylated cytosines, while unmodified cytosines are shown in grey. (**A**) Schematic structure of mt-tRNA^Met^ and the position of m^5^C (green circle) in the anticodon loop and variable loop. (**B**) Summary of the BS RNA-Seq results for mt-tRNA^Met^ in WT and NSUN2 -/- mouse liver. (C) Heatmaps of BS RNA-Seq reads for mt-tRNA^Met^ for WT and NSUN2 -/- showing cytosines of individual reads (on y-axis). (**D**) Schematic structure of mt-tRNA^Leu(UUR)^, mt-tRNA^His^, mt-tRNA^Glu^, mt-tRNA^Asn^ and mt-tRNA^Tyr^ and the position of m^5^C (green circle). (**E**) Summary of the BS RNA-Seq results for mt-tRNA^Leu(UUR)^, mt-tRNA^His^, mt-tRNA^Glu^, mt-tRNA^Asn^ and mt-tRNA^Tyr^ of WT and NSUN2 -/- mouse liver tissue. (**F**) Schematic structure of mt-tRNA^Leu(CUN)^ and the position of m^5^C (green circle). (**G**) Summary of the BS RNA-Seq results for mt-tRNA^Leu(CUN)^ of WT and NSUN2 -/- mouse liver tissue. (**H**) Schematic structure of mt-tRNA^Ser(AGY)^ and the position of m^5^C (green circle) in V-loop. (**I**) Summary of the BS RNA-Seq results for for mt-tRNA^Ser(AGY)^ in WT and NSUN2 -/- mouse liver tissue.

### Human NSUN2 methylates mitochondrial tRNAs at positions 48-50

Next, we intended to verify whether human NSUN2 is also responsible for methylation of mt-tRNA. To this end we performed targeted BS RNA-Seq using RNA isolated from primary fibroblasts from a patient harbouring a homozygous splice mutation in the *NSUN2* gene and that lacks a functional NSUN2 protein (NSUN2 mut/mut) (35). We used RNA from a healthy control (WT) and fibroblasts from a patient carrying loss-of-function mutations in *NSUN3* (NSUN3 mut/mut) (13) as controls. We analysed six mt-tRNAs that contain at least one cytosine in the V-loop region, namely mt-tRNA^Leu(UUR)^ (C48), mt-tRNA^His^ (C48), mt-tRNA^Glu^ (C49), mt-tRNA^Ser(AGY)^ (C48, C49 and C50), mt-tRNA^Tyr^ (C48) and mt-tRNA^Phe^ (C48) using BS RNA-Seq. This experiment revealed that m^5^C in the V-loop region of all six was decreased to background levels in NSUN2 mutant fibroblasts, but remained unaffected in the controls (**Figure 5A-D**). Of note, similarly to bovine mt-tRNA^Ser(AGY)^ the human counterpart contains three cytosines in the V-loop region (C48-50) (**Figure 5C-D**). However, in contrast to bovine mt-tRNA^Ser(AGY)^ which contains only m^5^C49 (3), in human mt-tRNA^Ser(AGY)^ all three positions are modified by NSUN2. None of the V-loop positions in human mt-tRNA were dependent on NSUN3 previously shown to introduce m^5^C34 in human mt-tRNA^Met^, indicative for non-redundant roles for these mitochondrial tRNA m^5^C methyltransferases (**Figure 5E-F**).

**Figure 5.**
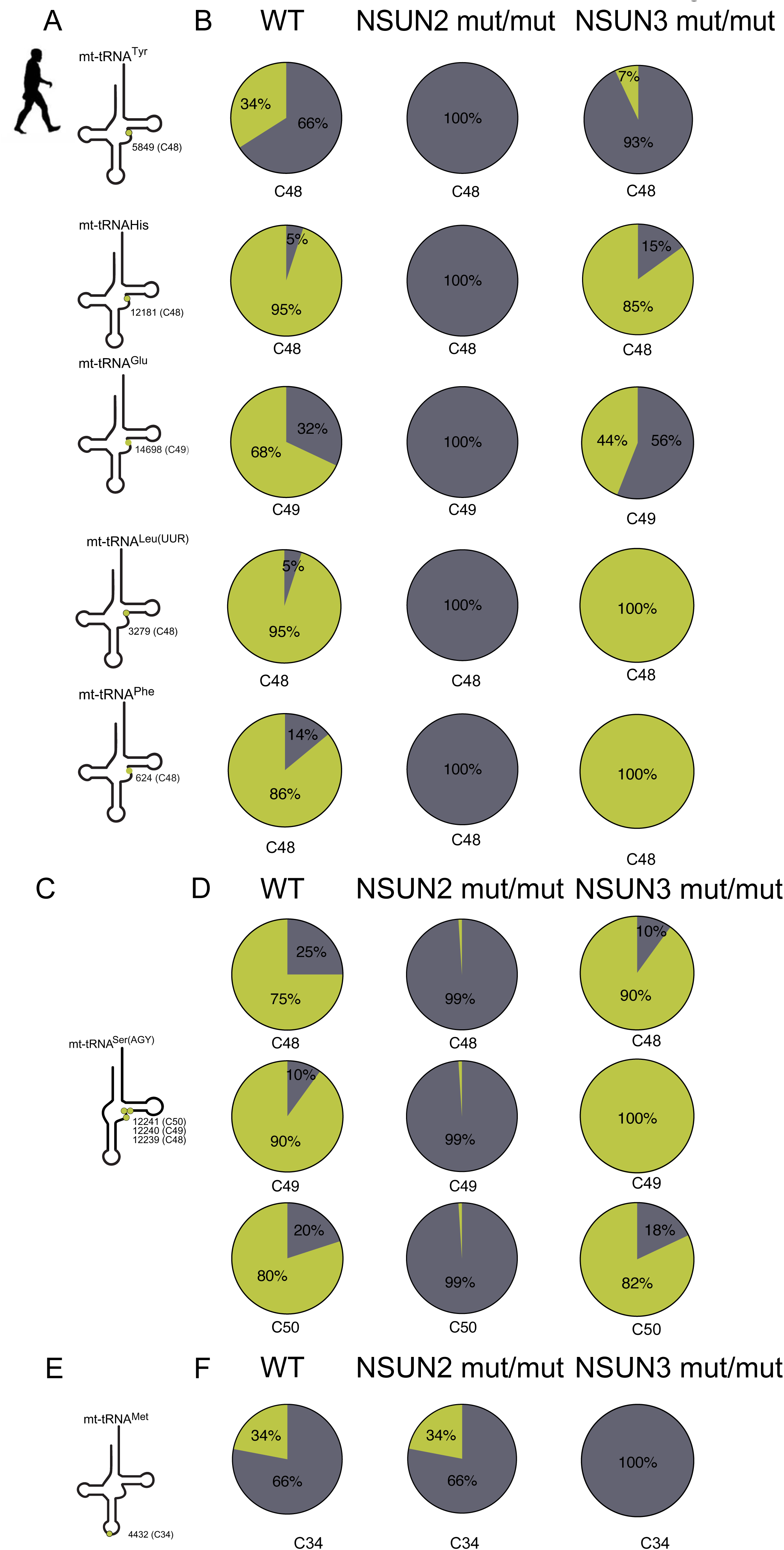
Human NSUN2 methylates mitochondrial tRNA postions 48-50. (**A-F**) Cytosine methylation of mt-tRNA of human fibroblasts. (**A**) Schematic structure of mt-tRNA^Tyr^, mt-tRNA^His^, mt-tRNA^Glu^, mt-tRNA^Leu(UUR)^ and mt-tRNA^Phe^. The green circles depict the positions of m^5^C in V-loop. (**B**) Summary of the BS RNA-Seq results for WT, NSUN2-null (NSUN2 mut/mut) and NSUN3-null (NSUN3 mut/mut) patient fibroblasts. (**C**) Schematic structure of mt-tRNA^Ser(AGY)^ and the positions of m^5^C (green circles) in the variable loop. (D) Summary of the BS RNA-Seq results for mt-tRNA^Met^ for WT fibroblasts, NSUN2 mut/mut and NSUN3 mut/mut. (**E**) Schematic structure of mt-tRNA^Met^ and the position of m^5^C (green circle) at position 34. (**F**) Summary of the BS RNA-Seq results for mt-tRNA^Met^ for WT fibroblasts, NSUN2 mut/mut and NSUN3 mut/mut.

To corroborate these results, we have generated an NSUN2 CRISPR/Cas9 knock-out cell line using human U-2 osteosarcoma (U2OS) cell line (NSUN2 KO) (**Supplementary Figure S1**). Firstly, we confirmed the lack of detectable m^5^C in the mt-tRNA V-loop region of mt-tRNA^Leu(UUR)^ (C48), mt-tRNA^His^ (C48), and mt-tRNA^Ser(AGY)^ (C48, C49 and C50) in the knock-out cells (**Figure 6**). Next, we set out to complement the NSUN2 defect by re-expressing the Flag-tagged version of the NSUN2 (NSUN2 KO + NSUN2-FLAG) that partially localises into human mitochondria (**Figure 2**). Transient expression of NSUN2.FLAG.STREP2 in the NSUN2 KO human cells resulted in partial restoration of m^5^C in mt-tRNA (**Figure 6A-C**). These results support the dependency of the mt-tRNA V-loop region m^5^C on NSUN2 and, importantly, strengthens the notion that the NSUN2 isoform encoded by the main splice mRNA splice variant (GenBank: NM_017755.6) is localised inside human mitochondria. In summary, our data show that inactivation of NSUN2 in mammalian species (mouse and man) results in the loss of the m^5^C modification in the V-loop region of mt-tRNAs, consistent with the mitochondrial localisation of a pool of the NSUN2 protein.

**Figure 6.**
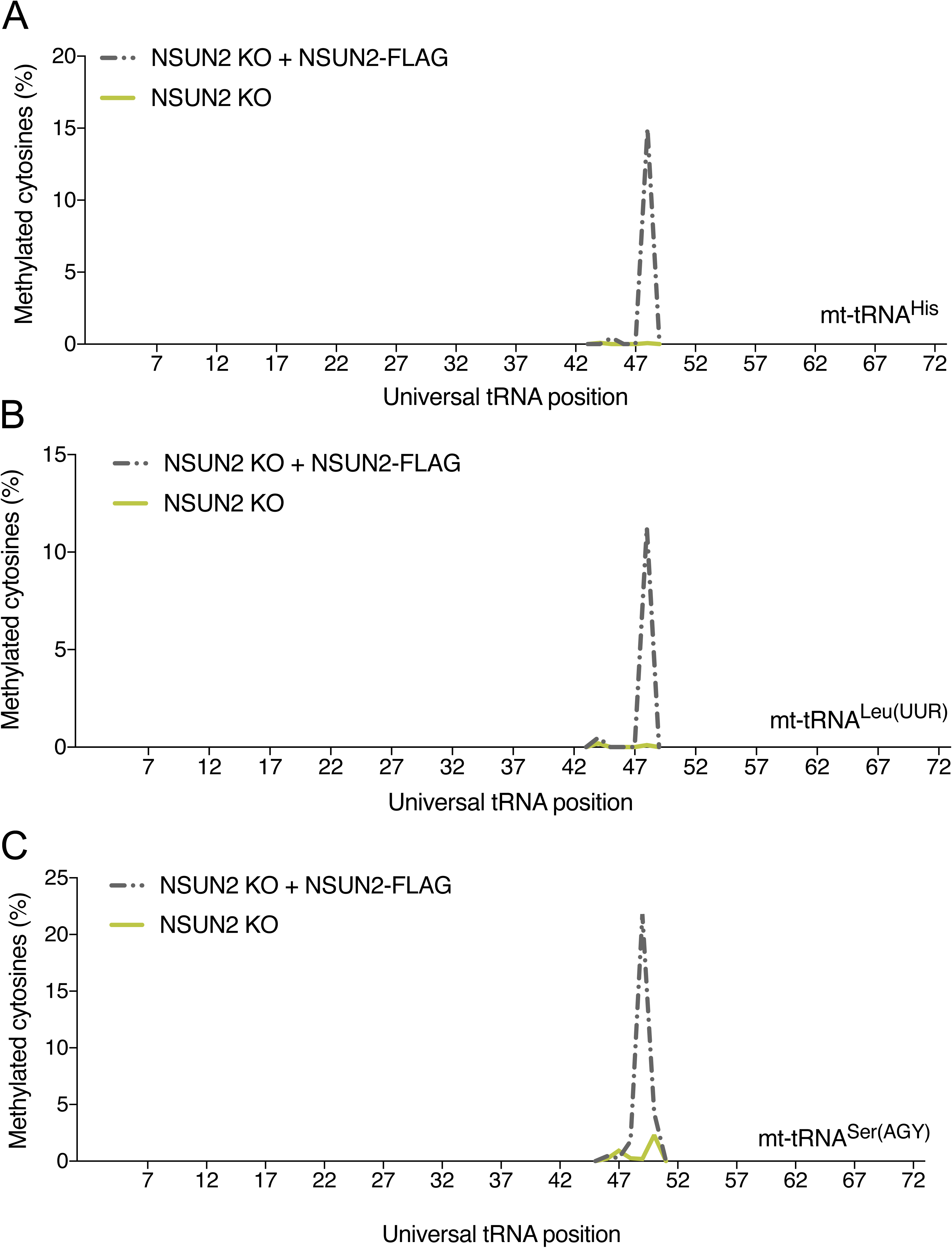
Transient expression of NSUN2 partially restores m^5^C in mt-tRNA. (**A-C**) Targeted BS RNA-Seq after transient expression of a NSUN2.FLAG.STREP2 construct in human NSUN2 KO cells. (A) Methylated cytosines of mt-tRNA^His^. (**B**) Methylated cytosines of mt-tRNA^Leu(UUR)^. (**C**) Methylated cytosines of mt-tRNA^Ser(AGY)^. The methylation levels have been adjusted to the percentage of the NSUN2.FLAG.STREP2 transfected cell (**Supplementary Figure S2**).

### Mitochondrial function upon inactivation of NSUN2

In the next set of experiments, we asked whether ablation of NSUN2 affects mitochondrial function *in vivo* in a mouse model through its impact on mitochondrial gene expression. NSUN2 knock-out mice are viable, but appreciably smaller than their wild-type and heterozygous littermates, with males being sterile and both genders showing cyclic alopecia (24) and brain development disorders (36). Firstly, we checked whether the absence of NSUN2 in mice affects mtDNA copy number. We analysed mtDNA copy number by qPCR in liver and heart tissue from NSUN2 knock out (NSUN2 -/-) mice and age-matched wild type control tissue, however, no significant difference was measured (**Figure 7A**). In order to investigate the effect of the lack of NSUN2 on the stability of mt-tRNAs, we performed northern blot analysis using mouse RNA isolated from liver and heart (**Figure 7B-C**). We found no significant differences in the steady-state levels of mouse mitochondrial NSUN2-targets (mt-tRNA^Glu^, mt-tRNA^His^ or mt-tRNA^Ser(AGY)^) as compared to mt-tRNA^Lys^, which has no cytosine in the V-loop region, hence is not a substrate of NSUN2. These results indicate that methylation of the V-loop region in mt-tRNAs by NSUN2 is not crucial for the stability of the mt-tRNAs, consistent with the existing data for cyto-tRNAs. Despite the normal steady-state levels of mt-RNA in NSUN2 -/- animals, their function could be perturbed by the lack of m^5^C in the V-loop region impacting intra-mitochondrial translation and consequently OXPHOS complex assembly and function. To address this, we analysed a variety of proteins of the OXPHOS system in the NSUN2 -/- mice by western blot. We compared heart and liver tissue of NSUN2 -/- animals with wild type mice and did not detect any appreciable differences in steady-state levels of OXPHOS complex proteins (**Figure 7D**). Taken together, our data indicate that the lack of m^5^C in the V-loop region does not have any profound effect on mitochondrial function.

**Figure 7.**
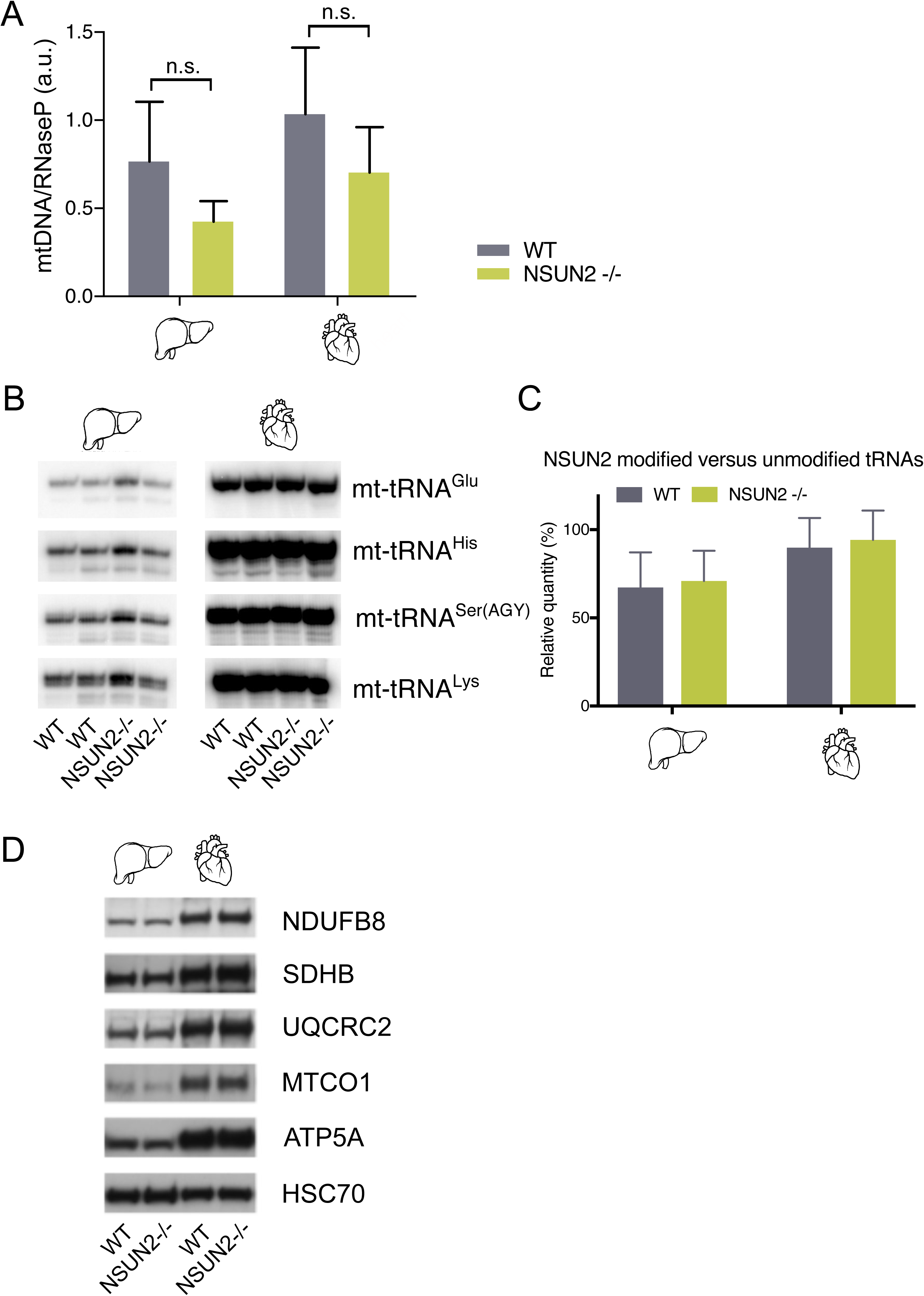
Mitochondrial function upon inactivation of NSUN2. (**A**) mtDNA copy number determination by qPCR of mtDNA relative to the nuclear RNase P gene in liver and heart of WT and NSUN2 -/- mice. qPCR was performed in triplicate and error bars indicate the s.d. of the mean. (**B**) Northern blot analysis of three mt-tRNAs that are modified by NSUN2 (mt-tRNA^Glu^, mt-tRNA^His^ and mt-tRNA^Ser(AGY)^ in WT and NSUN2 -/- liver and heart tissue. Mt-tRNA^Lys^, which is not methylated by NSUN2, is used as a loading control. (**C**) Quantification of the band intensities shown in B. (**D**) Western blot analysis of liver and heart for NDUFB8, SDHB, UQCRC2, MTCO1, ATP5A. HSC70 was used as loading control.

## Discussion

Recent years have witnessed a rapid improvement in our understanding of how the expression of mtDNA-encoded genes is regulated both in human health and in disease states (1,2). Dysfunction of mitochondrial gene expression, caused by mutations in either the mitochondrial or nuclear genomes, is associated with a diverse group of human disorders characterized by impaired mitochondrial respiration. Within this group, an increasing number of mutations have been identified in nuclear genes involved in mtRNA epitranscriptome shaping (13,37-46). Furthermore, there is increasing evidence that some of the most common heteroplasmic pathogenic mtDNA mutations perturb the introduction of mt-tRNA post-transcriptional modifications (14,47,48). Lastly, a number of mtRNA modifying enzymes have been identified and characterised using basic molecular and systems biology approaches (23,49-53). Despite this recent progress several mitochondrial RNA modifications, including m^5^C, await the responsible enzyme to be identified.

The NSUN2 protein has been reported to introduce m^5^C in several nuclear-encoded transcripts, including the V-loop and wobble position of tRNA, mRNAs, microRNAs and vault noncoding RNAs. In this work we showed mitochondrial localisation of NSUN2. Given its wide range of targets in cytoplasmic RNA, we investigated whether NSUN2 also methylates mtRNA. Our experimental work has revealed several NSUN2-dependent targets in mtDNA-encoded mammalian tRNAs localised in the V-loop region (positions 48, 49, 50). The m^5^C modification is most often found at positions 48 and 49 of the V-loop, located between the anticodon arm and the T-arm. Methylation at these positions on the nucleobase does not directly interfere with Watson-Crick base pairing, however its presence has been demonstrated to contribute to tRNA stability and protein synthesis (18). C48 has been shown to contribute to tRNA folding through the formation of a reverse Watson-Crick base pair with G15 in the D-loop, with one of these residues often modified to further stabilize the interaction (54). In the case of m^5^C, the contribution of the methyl group is not yet fully understood, but it may aid base stacking through increasing the hydrophobicity of the nucleobase, and/or promote Mg^2+^ binding which in turn increases stability (55). Previously published data showed that the stability of mammalian cytoplasmic decreases when NSUN2 and Dnmt2 are absent in mice (18,56). However, there is no evidence that loss of NSUN2 alone reduces the abundance of these cyto-tRNAs (36). Our analysis of mt-tRNAs modified by NSUN2 in the tissues of the NSUN2 -/- mouse also did not reveal any substantial changes in their stability, suggesting that similar to cyto-tRNAs, the lack of the V-loop m^5^C alone is not sufficient to destabilise tRNA. More detailed analysis will be required to establish the mitochondrial function of NSUN2. However, it is worth noting that the mechanism of how V-loop nucleotides contribute to mt-tRNA folding may be different as compared to cyto-tRNA. Enzymatic and chemical probing of bovine mt-tRNA^Phe^ have suggested that these tRNAs primarily rely on interactions between the D-stem and V-loop in order to achieve the L-shaped tertiary structure required (57), as the majority of mammalian mt-tRNAs lack the conserved residues, particularly G18, G19, Ψ55, and C56, that are required for the stabilising D-loop/T-loop interaction in cyto-tRNA.

Mutations in the NSUN2 gene have been associated with neurodevelopmental disorders in humans. Martinez *et al*. (35) and Abbasi-Moheb *et al*. (58) reported several families harbouring homozygous splice or stop mutations in *NSUN2* presenting with Dubowitz-like syndrome characterised by the constellation of mild microcephaly, intellectual disability, growth retardation, facial dysmorphism and muscular hypertonia. Furthermore, Khan *et al*. (59) reported a family with three affected children carrying a homozygous missense variant causing a mutation at a conserved residue (p.Gly679Arg) presenting with spasticity, ataxic gait, increase of liver enzymes and creatine phosphokinase in addition to dysmorphisms and developmental delay. Although the overall phenotype of the Dubowitz-like syndrome patients reported by Martinez et al. and Abbasi-Moheb et al. cannot be specifically linked to mitochondrial disorders, brain abnormalities, including microcephaly, have been commonly reported in *bone fide* mitochondrial respiratory chain disorders (13,60-62). Furthermore, the phenotypic features found in the family harbouring the missense *NSUN2* variant reported by Khan *et al*. could be considered as typical for mitochondrial disease. Careful follow up studies of the mitochondrial function (e.g. plasma lactate levels or TCA intermediates) in the reported and new cases could be considered in the view of our discovery of a mitochondrial function for NSUN2.

Our results imply that the lack of m^5^C in the V-loop region due to ablation of NSUN2 does not have any profound effect on mitochondrial function in differentiated cells. Based on previous studies in a mouse model, NSUN2 has been implicated in skin, germ cell and neural differentiation (24,36,63,64). The mitochondrial OXPHOS is crucial in the differentiation process (65,66). We speculate that in patients that carry a mutation in the *NSUN2* gene or in NSUN2 mouse knockout models, the aberrant modification of mt-tRNA could affect translation of mitochondrially-encoded proteins, perturbing the energy metabolism needed in differentiating cells. Therefore, future research on the mitochondrial function of NSUN2 could focus on early stages of neural development.

## Acknowledgements

We would like to thank the members of the Mitochondrial Genetics Group at the MRC-MBU for stimulating discussions during the course of this work.

## Authors’ contribution

L.V.H. conceived, designed and performed the experiments, undertook the data analysis and wrote the manuscript; S-Y. L, S.S., J-S. K. and H-W. R. contributed the proximity labelling analysis and generated Figure 1; B.J.M performed immunostaining experiments (Figure 2A-B); C.A.P. contributed to the northern blotting analysis; C.G. analysed the patient related data; M.F. provided the NSUN2 mouse model; J.G.G. provided the NSUN2 patient cells; D.B. and E.M. provided the CRISPR/Cas9 NSUN2 KO cells; M.M. conceived and oversaw the project and wrote the manuscript. All authors commented and approved the final version of the manuscript.

