## Supplementary material for "NSUN2 introduces 5-methylcytosines in mammalian mitochondrial tRNAs"

Lindsey Van Haute *et al*.

CONTENTS

Supplementary Figure S1 | Generation and characterisation of NSUN2 CRISPR/Cas9 KO cells – page 2

Supplementary Figure S2 | Transient expression of NSUN2 in human NSUN2 KO cells – page 3

Supplementary Table S1 | List and sequences of gRNA used to generate human NSUN2 KO line – page 4

Supplementary Table S2 | List of oligonucleotides used in this study

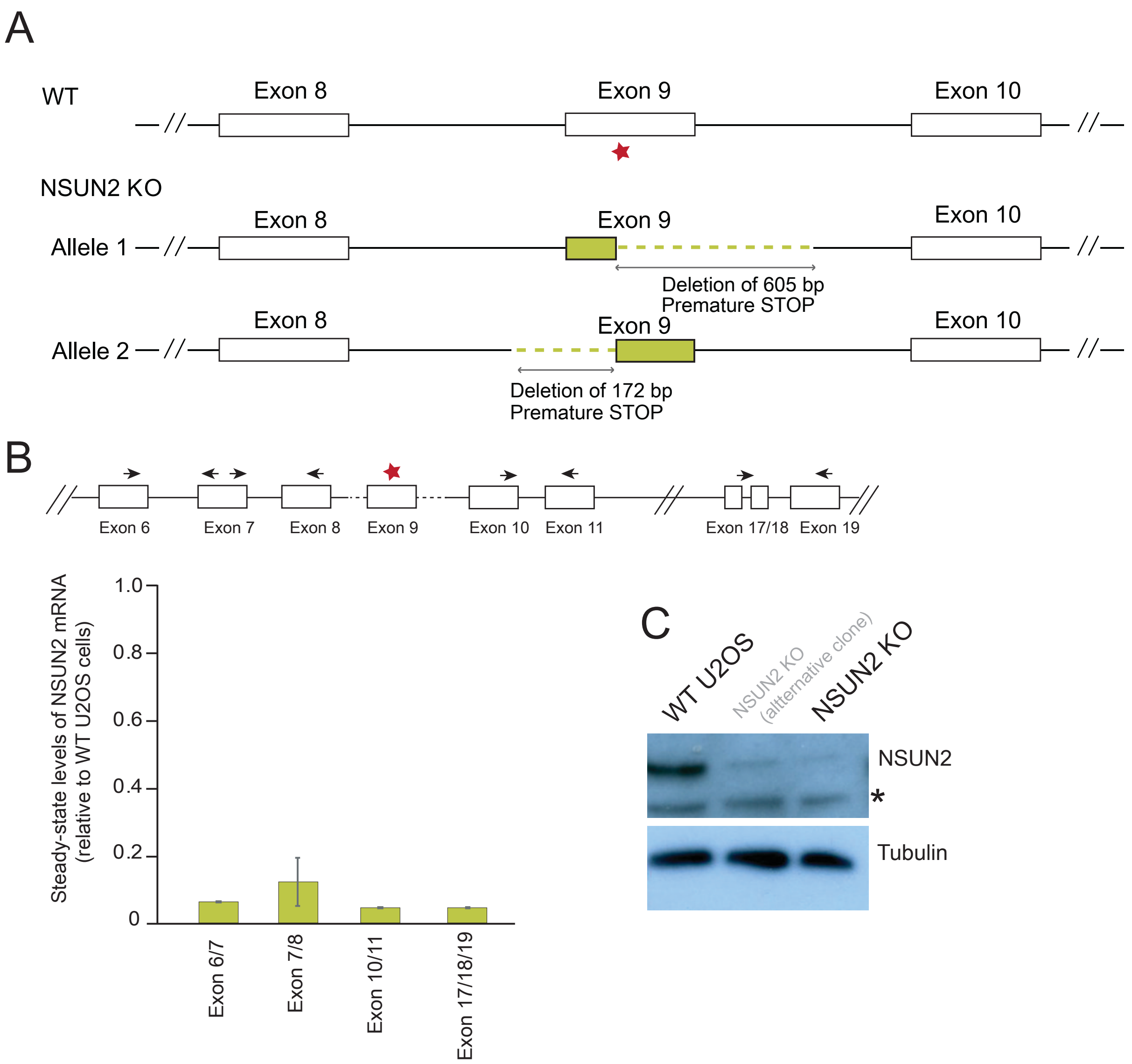


**Supplementary Figure S1 |** Generation and characterisation of NSUN2 CRISPR/Cas9 KO cells
(**A**) Schematic representation of a fragment of the NSUN2 gene structure spanning exons 8 to 10. Red star indicates the site in exon 9, which was targeted by the CRISPR/Cas9 gRNAs (**Supplementary Table S1**). Genomic changes in the U2OS NSUN2 KO cell line as detected by PCR and Sanger sequencing are indicated in green and described. (**B**) *Top*: the NSUN2 gene structure spanning exons 6 to 19 with indicated RT-qPCR primer binding sites. *Bottom*: RT-qPCR analysis of NSUN2 mRNA expression levels in the NSUN2 KO cells for 4 pairs of primers. The expression was normalised to GAPDH. n=2. (**C**) Western blot analysis of the steady-state levels of NSUN2 in U2OS WT and NSUN3 KO cell line. Note: alternative CRISPR/Cas9 KO line is shown, which was not used in the present study.

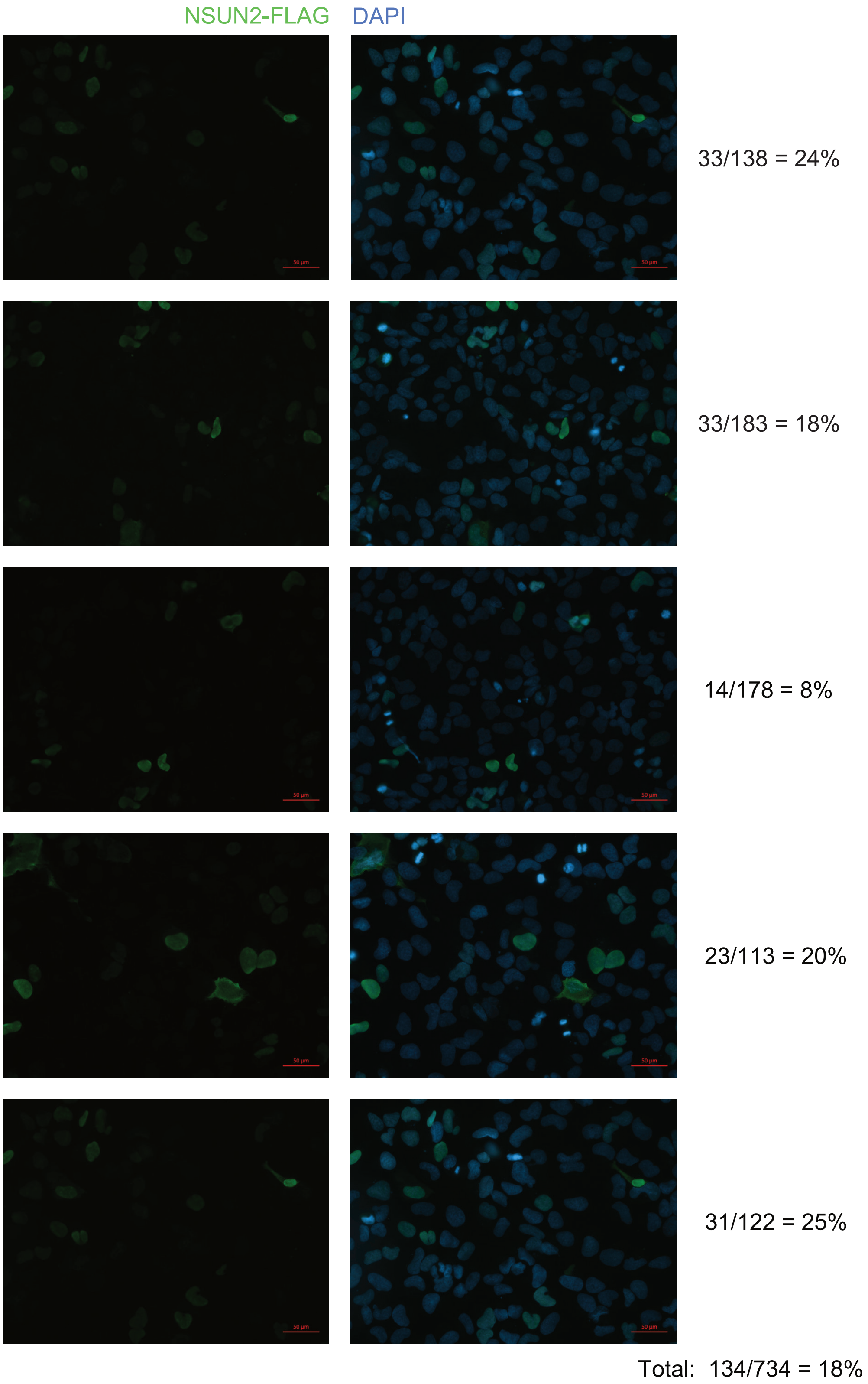


**Supplementary Figure S2 |** Transient expression of NSUN2 in human NSUN2 KO cells

Representative images of immunofluorescence labelling after transient expression of a NSUN2.FLAG.STREP2 construct in human NSUN2 KO cells. Cells were stained for FLAG (green) and DAPI (blue). Scale bar: 50μM. Average percentage of transfected cells was 18 %.

**Supplementary Table S1** | List and sequences of gRNA used to generate human NSUN2 KO line

| **gRNA name** | **gRNA sequence** |
|---|---|
| gRNA_C321A_1_fwd | pACCGCAATCCGCAGCTGTAAGCTA |
| gRNA_C321A_1_rev | pAAACTAGCTTACAGCTGCGGATTG |
| gRNA_C321A_2_fwd | pACCGGTGTTCACTAAACCCTATTG |
| gRNA_C321A_2_rev | pAAACCAATAGGGTTTAGTGAACAC |

**Supplementary Table S2 |** List of oligonucleotides used in this study

| Purpose | | 5' to 3' sequence |
|---|---|---|
| **Primers used for cloning into pcDNA5-FST2** | | |
| NSUN2 start | forward | GACGGTACCATGGGGCGGCGGTCGCGGGG |
| NSUN2 stop | reverse | GTCCTCGAGCCGGGGTGGATGGACCCCC |
| **qPCR primers for DNA analysis** | | |
| mt-CO1 | forward | TGCTAGCCGCAGGCATTACT |
| | reverse | CGGGATCAAAGAAAGTTGTGTTT |
| RNaseP | forward | GCCTACACTGGAGTCCGTGCTACT |
| | reverse | CTGACCACACACGAGCTGGTAGAA |
| **qPCR primers for RT-qPCR** | | |
| NSUN2 ex17-18 | forward | AAGCAAAGGACCTGGCAAAG |
| NSUN2 ex19 | reverse | CAGCCCCATCATCCTGAGAT |
| NSUN2 ex10 | forward | AGTGGATGCCTGGAATCACA |
| NSUN2 ex11 | reverse | GGGAACATGGTAGGTCGGAT |
| NSUN2 ex6 | forward | CTGGCTCAAAGACCACACAG |
| NSUN2 ex7 | reverse | GTTGACCACCATGATGCAGG |
| NSUN2 ex7 | forward | CCTGCATCATGGTGGTCAAC |
| NSUN2 ex8 | reverse | TCTCATAGTGCCGTCTCCAC |

**Supplementary Table S2 - continued**

| Targeted RNA BS-seq primers human | | |
|---|---|---|
| hMT-TS2 RT primer | | TAAAAAAACCATATTATTAAACA |
| hMT-TS2 1stage PCR primer with overhang | forward | TCGTCGGCAGCGTCAGATGTGTATAAGAGACAGGTTTATAAGAATTGTTAATTTATG |
| hMT-TS2 1stage PCR primer with overhang | reverse | GTCTCGTGGGCTCGGAGATGTGTATAAGAGACAGTAAAAAAACCATATTATTAAACA |
| hMT-TM RT primer | | TTAGTTAAATAAGTTATTGGGT |
| hMT-TM 1stage PCR primer with overhang | forward | TCGTCGGCAGCGTCAGATGTGTATAAGAGACAGTTAGTTAAATAAGTTATTGGGT |
| hMT-TM 1stage PCR primer with overhang | reverse | GTCTCGTGGGCTCGGAGATGTGTATAAGAGACAGAACCAACATTTTCAAAATA |
| hMT-TH RT primer | | TAAAAATCATAAACCTC |
| hMT-TH 1stage PCR primer with overhang | forward | TCGTCGGCAGCGTCAGATGTGTATAAGAGACAGTTAAAATATTAGATTGTGAATTTG |
| hMT-TH 1stage PCR primer with overhang | reverse | GTCTCGTGGGCTCGGAGATGTGTATAAGAGACAGAATAAAAAATCATAAACCTC |
| hMT-TL1 RT primer | | TATTAAAAAAAAAAAATTAAACCTC |
| hMT-TL1 1stage PCR primer with overhang | forward | TCGTCGGCAGCGTCAGATGTGTATAAGAGACAGGGTAATTGTATAAAATTTAAAAT |
| hMT-TL1 1stage PCR primer with overhang | reverse | GTCTCGTGGGCTCGGAGATGTGTATAAGAGACAGTATTAAAAAAAAAAAATTAAACCTC |
| hMT-TE RT primer | | TATTCTCACACAAACTACAACCA |
| hMT-TE 1stage PCR primer with overhang | forward | TCGTCGGCAGCGTCAGATGTGTATAAGAGACAGAATATAATGATGGTTTTTTATA |
| hMT-TE 1stage PCR primer with overhang | reverse | GTCTCGTGGGCTCGGAGATGTGTATAAGAGACAGTATTCTCACACAAACTACAACCA |
| hMT-TF RT primer | | TATTTATAAAATAATATAAACC |
| hMT-TF 1stage PCR primer with overhang | forward | TCGTCGGCAGCGTCAGATGTGTATAAGAGACAGTAAAGTAATATATTGAAAATGTTT |
| hMT-TF 1stage PCR primer with overhang | reverse | GTCTCGTGGGCTCGGAGATGTGTATAAGAGACAGTATTTATAAAATAATATAAACC |
| hMT-TY RT primer | | ATAATAAAAAAAAACCTAACCCC |
| hMT-TY 1stage PCR primer with overhang | forward | TCGTCGGCAGCGTCAGATGTGTATAAGAGACAGGTTGAGTGAAGTATTGGATTGTAA |
| hMT-TY 1stage PCR primer with overhang | reverse | GTCTCGTGGGCTCGGAGATGTGTATAAGAGACAGATAATAAAAAAAAACCTAACCCC |

**Supplementary Table S2 - continued**

| Targeted RNA BS-seq primers mouse | | |
|---|---|---|
| mMT-TH RT primer | | AATAAATAAAAAAATTTATTTCC |
| mMT-TH 1stage PCR primer with overhang | forward | TCGTCGGCAGCGTCAGATGTGTATAAGAGACAGAAAATATTAGATTGTGAATTTG |
| mMT-TH 1stage PCR primer with overhang | reverse | GTCTCGTGGGCTCGGAGATGTGTATAAGAGACAGAATAAATAAAAAAATTTATTTCC |
| mMT-TL1 RT primer | | TATTAAAAAGAAAATTTAAACCTC |
| mMT-TL1 1stage PCR primer with overhang | forward | TCGTCGGCAGCGTCAGATGTGTATAAGAGACAGTGTGTAAGATTTAAAATTTTGT |
| mMT-TL1 1stage PCR primer with overhang | reverse | GTCTCGTGGGCTCGGAGATGTGTATAAGAGACAGTATTAAAAAGAAAATTTAAACCTC |
| mMT-TL2 RT primer | | TACTTTTATTTAAATTTACACCA |
| mMT-TL2 1 stage PCR primer with overhang | forward | TCGTCGGCAGCGTCAGATGTGTATAAGAGACAGATAATAGTAATTTATTGGTTTTAGGA |
| mMT-TL2 1stage PCR primer with overhang | reverse | GTCTCGTGGGCTCGGAGATGTGTATAAGAGACAGTACTTTTATTTAAATTTACACCA |
| mMT-TY RT primer | | TAATAAAAAAAAAATTTAAACCTC |
| mMT-TY 1 stage PCR primer with overhang | forward | TCGTCGGCAGCGTCAGATGTGTATAAGAGACAGGAGTAAGTATTAGATTGTAAAT |
| mMT-TY 1 stage PCR primer with overhang | reverse | GTCTCGTGGGCTCGGAGATGTGTATAAGAGACAGTAATAAAAAAAAAATTTAAACCTC |
| mMT-TN RT primer | | CTAAATTAACAAAAATTAAACCTA |
| mMT-TN 1 stage PCR primer with overhang | forward | TCGTCGGCAGCGTCAGATGTGTATAAGAGACAGTAATAGGGTATTTAGTTGTTAA |
| mMT-TN 1 stage PCR primer with overhang | reverse | GTCTCGTGGGCTCGGAGATGTGTATAAGAGACAGCTAAATTAACAAAAATTAAACCTA |
| mMT-TS2 RT primer | | TAAAAAAACCATATTTTTAAACA |
| mMT-TS2 1stage PCR primer with overhang | forward | TCGTCGGCAGCGTCAGATGTGTATAAGAGACAGTTGTAAGAATTGTTAATTTATG |
| mMT-TS2 1stage PCR primer with overhang | reverse | GTCTCGTGGGCTCGGAGATGTGTATAAGAGACAGTAAAAAAACCATATTTTTAAACA |
| mMT-TE RT primer | | TATTTCTACACAACATTCAACTA |
| mMT-TE 1stage PCR primer with overhang | forward | TCGTCGGCAGCGTCAGATGTGTATAAGAGACAGTGATGATTTTTTATGTTATTGG |
| mMT-TE 1stage PCR primer with overhang | reverse | GTCTCGTGGGCTCGGAGATGTGTATAAGAGACAGTATTTCTACACAACATTCAACTA |
